# Diversity, abundance, and host specificity of the human skin associated circular and single stranded DNA virome

**DOI:** 10.1101/2022.05.22.492996

**Authors:** Ema H. Graham, Michael S. Adamowicz, Peter C. Angeletti, Jennifer L. Clarke, Samodha C. Fernando, Joshua R. Herr

## Abstract

The human skin is our point of contact with the microbial world, yet little is known about the diversity of the skin virome. Studies of the human skin virome have focused on bacteriophage and double-stranded DNA viral genomes, however, there have been few efforts to characterize circular single-stranded DNA viruses that populate human skin. Here, we evaluate the diversity of the circular single-stranded DNA virome collected across three anatomical skin locations from 60 human individuals with five time-point collections spanning six-months. Our analyses resulted in the identification of 272 novel and unique Rep-encoded single-stranded DNA viruses associated with human skin. Sequence similarity networks and maximum likelihood estimations of the Rep and Capsid protein amino acid sequences from our sequencing and public database references, reveal family level stability of the *Cressdnaviricota* across the study participants and a larger host-range than previously thought for these putative multi-host pathogens.

## INTRODUCTION

Understanding viral diversity in the environment and the potential for epidemics driven by host switching is imperative to anticipate and combat the emergence of novel zoonotic viral pathogens. To develop effective pathogenic mitigation strategies, we first must be able to identify viral communities and predict those that have the capability to cause significant pathogenesis to humans after acquisition from non-human hosts, such as wild or domesticated animals. This presents a challenge since viral diversity in current reference databases is poorly characterized due to the limited number of studies investigating environmental, human, and animal associated viromes. Notably, metagenomic sequencing techniques have contributed to the discovery and identification of novel DNA and RNA viral genomes in the environment [1].

The human skin is the first point of contact with many environmental pathogens, such as viruses. Most zoonotic viral emergence studies have been directed at viral vectors in aerosols or through infections of the digestive tract and have consisted of the prediction of host switching events to predict pathogenesis in human hosts [2, 3]. However, few studies have directly addressed the presence of environmentally transmitted viruses on human skin and identified the potential to become human pathogens via previous host switching events.

Recent studies have addressed the composition of the human skin virome through metagenomic sequencing techniques [4–9]. The human skin virome is mainly composed of transient environmentally contracted viruses [5,6,9]. Studies addressing the diversity of the skin virome have almost exclusively utilized DNA sequencing techniques and have characterized the double-stranded DNA (dsDNA) virome, which mostly consists of bacterial infecting viruses, such as bacteriophage [5, 9]. The abundance of bacteriophage genomes identified in human skin virome studies is not surprising due to established commensal bacterial communities on the skin and the overabundance of *Caudovirales* bacteriophage genomes in viral reference databases [10–15]. Though these viruses may have secondary human health implications through the control of human skin bacterial microbiome population dynamics [5,8,16], most dsDNA skin associated viruses are not widely viewed as being directly detrimental to human health [7, 8].

Small circular DNA viruses, in the viral realm of *Monodnaviria*, on the human skin are diverse, abundant, and have largely been uncharacterized [4,5,9]. Arguably the most well studied of the circular DNA viruses are the commensal dsDNA skin opportunistic pathogens Papillomaviruses and Polyomaviruses, both of which have been associated with cancer and abnormal skin growth [4,17,18]. Excluding those groups, the vast amount of small circular single stranded DNA (ssDNA) viruses are novel and, therefore, our knowledge of their diversity is lacking [9, 19]. The extent to which the ssDNA skin viruses cause disease is not known.

One such diverse group of small circular ssDNA viruses is the *Cressdnaviricota*. This phylum is characterized by viruses with small genomes ranging from 0.98 to 6.30 kb in size that encode a unique rolling circle replication associated (Rep) protein [20]. Within the *Cressdnaviricota* phylum there are families known to infect eukaryotes including: plants (*Nanoviridae* and *Geminiviridae*), diatoms (*Bacilladnaviridae*), amoebas and protozoa (*Naryaviridae*, *Nenyaviridae*, and *Vilyaviridae*), fungi (*Genomoviridae*), humans and other animals (*Circoviridae*, *Genomoviridae*, *Smacoviridae*, and *Redondoviridae*) [20–26]. In addition to infecting eukaryotes, there is evidence to suggest that the *Smacoviridae*, a family of fecal associated Cressdnaviruses, broadly infect the Archaea [27]. Though Rep encoding viruses have been identified across a broad range of host organisms, the extent of their host associations is not understood. Due to the natural genomic variation of the *Cressdnaviricota*, it has been hypothesized that these viruses can broadly infect multiple different types of hosts and cell types [28]. Additionally, many of these viruses have been peripherally associated with acute respiratory distress, severe encephalitis, and systemic issues resulting in death as observed in both animals and humans [23, 29–31]. These observations make the *Cressdnaviricota* a prime candidate for studies focused on zoonotic and viral emergence in relation to human health.

We recently studied the human skin virome [9] and, in agreement with another study focused on animal skin virome diversity [32], we identified Cress-DNA-like viruses as being highly abundant. However, due to a lack of knowledge regarding the diversity of Cress-DNA-like viruses, we previously did not focus on identifying these ssDNA viruses past the phylum level of the *Cressdnaviricota* [9]. In the present study, we address the composition of novel ssDNA viruses on the human skin and characterize their diversity, stability, and their potential human health implications. First, we identified circular viral genomes from human skin metagenomes collected and sequenced from 60 individuals across three skin anatomical locations (left hand, right hand, and scalp) spanning five longitudinal samples taken across a six-month time-period post initial collection. Next, using data from metagenomic sequencing studies and public databases of viral diversity, we identified and predicted novel animal and environmentally contracted viral taxonomic groups that have been identified directly from the human skin, therefore eliminating the ambiguous question of transfer potential from animal vectors. Amino acid sequences of the Rep and Capsid genes encoded by both identified and novel skin associated Cressdnaviruses were then used to investigate phylogenetic classifications, host range, and ancestral host switching patterns in an attempt to predict potential animal host vectors.

## RESULTS

### Human skin associated viral metagenome assemblies

Human skin virome samples were collected from 60 subjects across three anatomical locations with five repeated collections performed over a six-month interval after the initial collection time point. Following DNA sequencing, viral metagenome assemblies from these samples were computationally assessed for presence of small circular DNA viruses. We assembled a total of 87,106 viral metagenome contigs from the skin samples of the 60 subjects in this study. A total of 62,101 viral metagenome contigs were greater than 1kb in size. Accounting for genomes found across multiple subjects, we identified a total of 683 unique viral genomes that were identified as having characteristics of complete small single-stranded circular viruses. These 638 viral genomes were further grouped based on their gene synteny and homology to known viral phylums or families. The groupings consisted of 272 *Rep* containing *Cressdnaviricota*, 2 *Anelloviridae*, 25 *Inoviridae* (22 of which contained a Zot cholera toxin gene), 27 *Papillomaviridae*, 2 *Polyomaviridae*, 67 *Microviridae*, 26 unclassified phage, and 93 viral contigs that remained unclassified but contained a capsid gene and exhibited homology to other small circular viruses (Fig. 1A).

**Fig 1.**
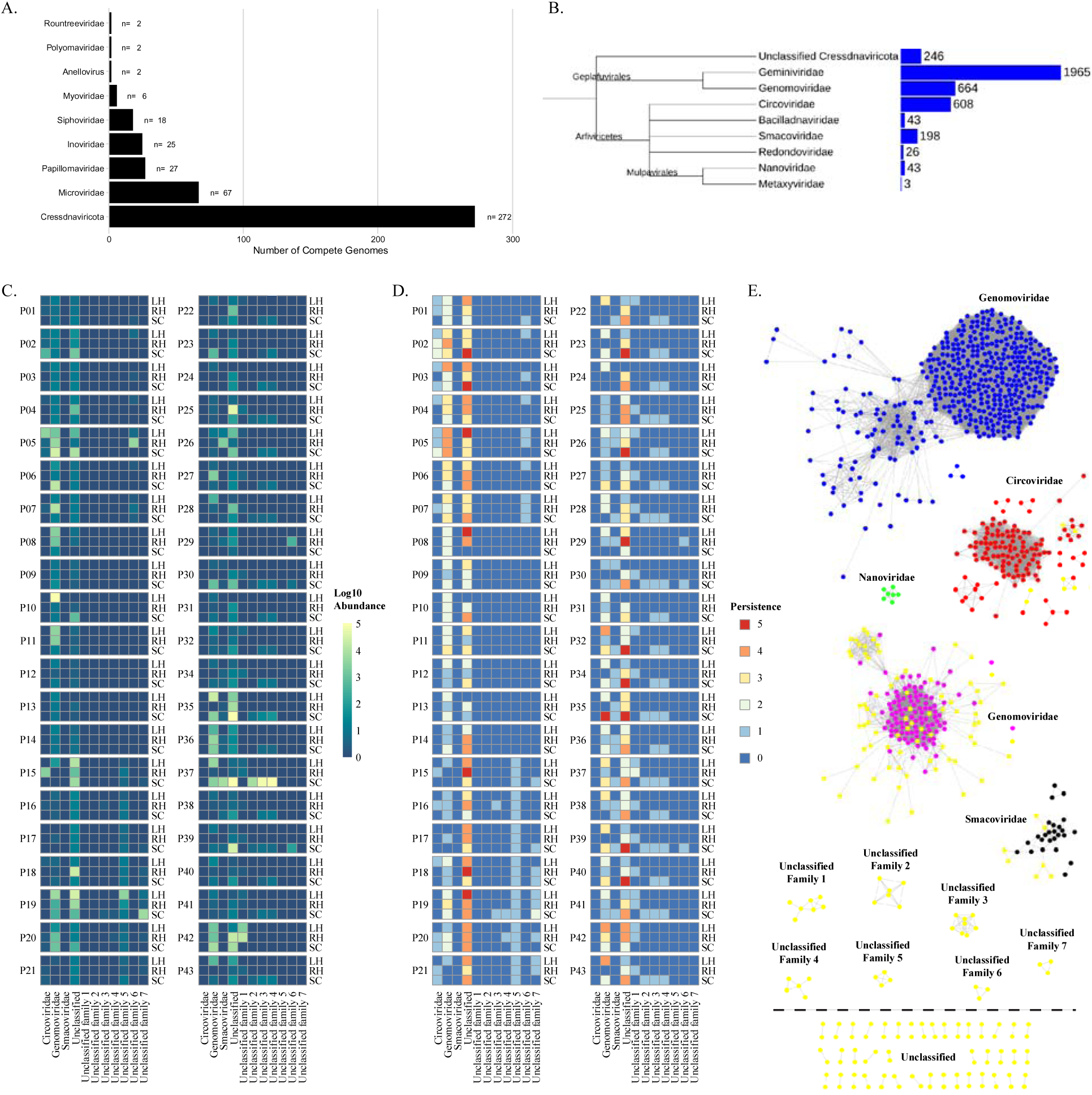
(A.) Distribution and classification of small circular DNA viruses identified from a meta-assembly and within subject assemblies from human skin associated viral metagenomic sequencing samples. (B.) Common tree of NCBI reference *Cressdnaviricota* family level phylogeny and the number of taxonomies associated with each family as reported by NCBI taxonomy [67]. Tree was generated using NCBI Taxonomy’s common tree tool [67] and visualized using iTOL [62]. Branch lengths are not representative of phylogenetic distance. *Cressdnaviricota* associated with the human skin diversity, abundance, and stability over time. (C.) Log10 total abundance summed across all time points for a subject at an anatomical location for the identified family level groupings of the novel human skin associated Cressdnaviruses. (D.) Persistence of the Cressdnavirus family level grouping of the novel human associated Cressdnaviruses across the five time points is represented by the color scale with red being present in five out of the five time points and blue being present in zero out of the five time points. For (C.) and (D.) heatmaps each row is an anatomical location (LH = Left Hand; RH = Right Hand; SC = Scalp). Rows are broken up by subject. (E.) Sequence similarity network of *Cressdnaviricota* reference sequences obtained from NCBI and the human skin associated novel CRESS-like viruses identified in this study. EFI-EST was used to conduct pairwise alignments of the conserved amino acid sequences of the *Cressdnaviricota* phylum Rep gene [55]. For family level clustering an E-value cutoff of 10^-60^ was used. Cytoscape was used for network visualization and color labeling [56].

### Identification of human skin associated CRESS-DNA genomes

We found complete viral genomes from the human skin viral metagenome samples that contained Rep genes, a unique characteristic of circular replication associated (Rep)-encoding single-stranded (CRESS) DNA viruses. These circular genomes were therefore classified as belonging to the viral phylum *Cressdnaviricota*. Due to Cressdnaviruses being poorly described in viral reference databases, many of these viral genomes had low percent identity when compared to known reference genomes using BLASTn or other viral annotation programs (e.g. Kaiju and Kraken2), making their taxonomic classification at levels below phylum level difficult and ambiguous (Fig. 1B) [33–35]. Therefore, to predict family level classification of these human skin associated assembled genomes, Rep gene translated amino acid sequence similarity networks were employed to take advantage of the conservation of the Rep gene sequence across *Cressdnaviricota* viral families (Fig. 1E). Using the amino acid sequence of the assembled viral genomes and a reference dataset obtained from NCBI containing all available Rep gene sequences with greater than 90% sequence similarity difference, sequence similarity networks were utilized to assess the taxonomic family level grouping of the 272 human skin associated *Cressdnaviricota* viruses identified in our dataset (Fig. 1E). For this initial family level classification assessment, we conducted reference sequence similarity network clustering using only full-length Rep amino acid sequences with known and reported family level classifications.

### Human skin associated *Cressdnaviricota* abundance and stability over time

The abundance (Fig. 1C) and persistence (Fig. 1D) over time (i.e., stability) of the putative family level groupings in our dataset were assessed. The family *Genomoviridae* did not only contain the most identified human skin associated viral genomes from our dataset as determined via SSN clustering, but they were also the most highly abundant of all the family level groupings present in our dataset, with the exception of family level clustering that remained unclassified. Additionally, the *Genomoviridae* were found to be stable and persistently present across at least four out of the five time points for many individual subjects and was found across multiple anatomical locations within an individual.

### Phylogenetic relationships among human skin associated and known human anatomical site associated *Genomoviridae*

Investigation of the human host associated *Genomoviridae* by anatomical site revealed a lack of body site affinity for the known human associated *Genomoviridae* Rep amino acid sequences (Fig. 2). This is indicative of *Genomoviridae* acting as a generalist pathogen since they did not discriminate across cell types. This is highlighted by regions of the phylogenetic tree where some clades are polytomies and are represented by being sourced from multiple human cell types. This can be observed most noticeably in the clade consisting of Rep protein sequences from *Genomoviridae* viruses isolated from human fecal samples (AJD20393.1, AJD20391.1), a throat swab (QCT85019.1), cerebral fluid (AJD20385.1, AJD20389.1, YP_00130661.1, AJD20387.1), cervix (QDF60272.1, QDF60273.1) and human skin virome samples from this study (CENOTE_1_10_META20738_2, CENOTE_META_662, CENOTE_META_662_2, CENOTE_1_10_META_20738). These clades of the *Genomoviridae* phylogeny are characterized by regions of high sequence similarity within these Rep genes. This observation suggests, in contrast with *Smacoviridae*, that the *Genomoviridae* are associated with numerous different human cell types and/or locations of isolation.

**Fig 2.**
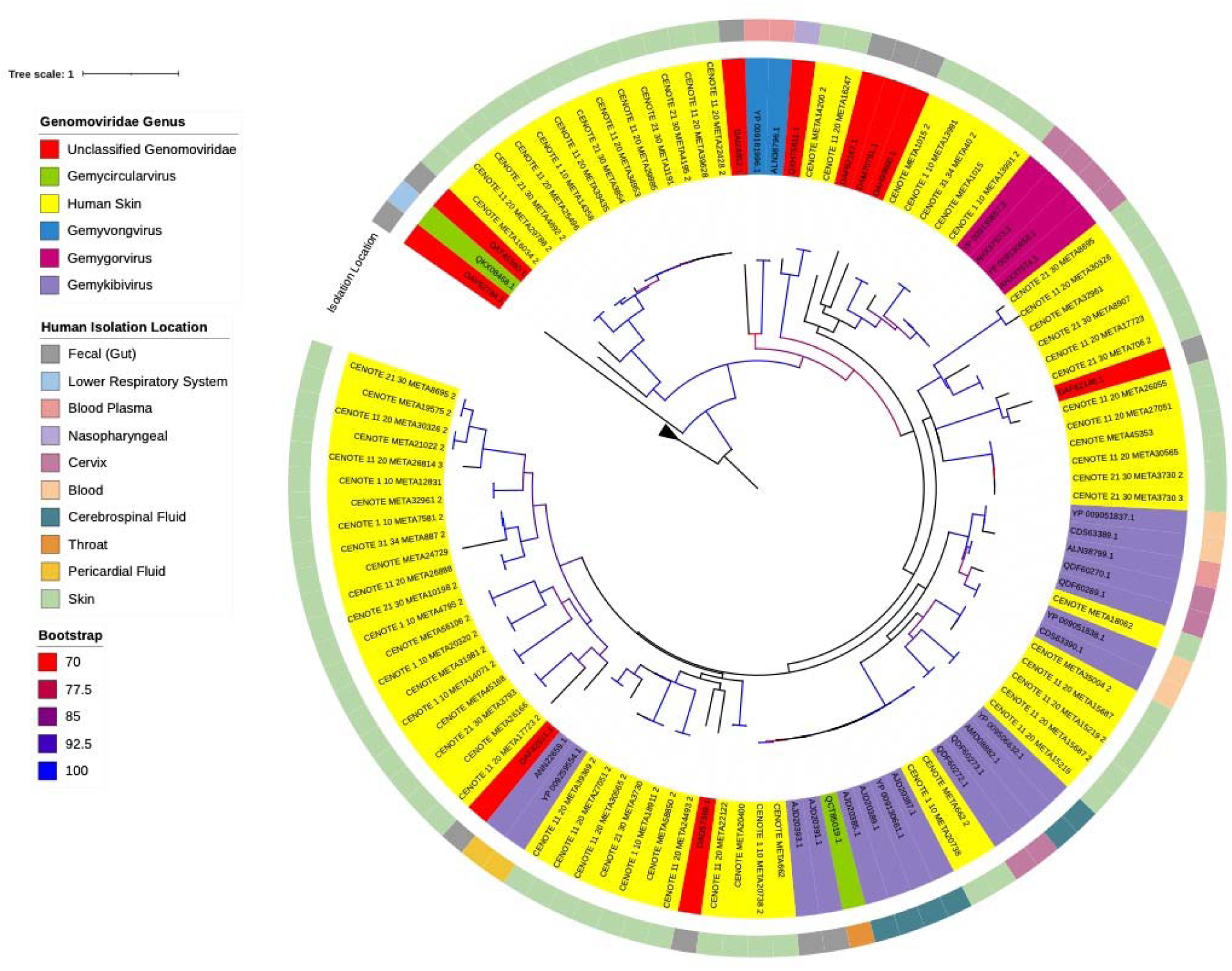
Maximum likelihood phylogenetic analysis of reference human associated *Genomoviridae* Rep protein amino acid sequences and identified novel human skin associated *Genomoviridae* Rep proteins. Reference sequences were downloaded from NCBI. Protein sequences were aligned using MAFFT inferred using IQTree with flags for ultrafast bootstrap determination and visualized using iTOL [59–62] using the best fit substitution model of VT+F+R4, as determined by IQTree. Branches with >70% bootstrap support are gradient colored with red representing 70% support and blue representing 100% support. Color coordination of genus classification (clade coloring) and reported human body sampling location (outermost ring) are represented in the figure legend. Four Rep gene amino acid sequences from differing Geminiviruses were used as an outgroup.

### Phylogenetic relationships among human skin associated and known human anatomical site associated *Smacoviridae*

Due to sequence conservation of the Rep gene [28, 36], all available full length Rep genes that were isolated from humans were used for phylogenetic assessment of the two identified family level groupings from our dataset. Skin associated *Cressdnaviricota* have not previously been isolated or investigated outside of reporting presence in diversity and meta-analysis studies of skin virome diversity [5, 9]. We wanted to determine if body site differences could be observed and to determine how closely these might relate to novel human skin associated *Cressdnaviricota* compared to previously discovered human associated and human pathogen associated *Smacoviridae* and *Genomoviridae*. Additionally, we wanted to investigate body site specific differences in the *Cressdnaviricota* to observe if human skin viruses shared homology and close common ancestors as they were isolated from the same host. By doing so, we reasoned that this may give insight into environmental contraction and transmission range of the two families of viruses, through direct touch, inhalation, and digestion.

All previously reported and published human associated *Smacoviridae* genomes were human gut associated and identified through metagenomic analysis of human fecal samples [30, 38]. Environmental genomes from the *Smacoviridae* have been identified from human wastewater and sewage systems, however, since they were not directly associated with samples collected directly from humans, we did not include these accessions in the phylogenetic analysis of total human associated *Smacoviridae*. Maximum likelihood phylogenies of the human skin associated and known human associated *Smacoviridae* Rep gene amino acid sequences exhibited strong clustering by human body site association (Fig. 3). The human skin *Smacoviridae* grouping was distinctly separate from that of all other human fecal associated *Smacoviridae* with the exception of one unclassified genome (DAL02127.1). Of the human associated *Smacoviridae* with a known sub-family classification, the human skin associated *Smacoviridae* likely shared a closer common ancestor to that of the *Porpismacoviridae*. Distant ancestry and the separate clustering of the human associated skin *Smacoviridae* and known human associated *Huchismacoviridae* was notable as the genus *Huchismacovirus* consists of the most abundantly identified human associated viral genomes.

**Fig 3.**
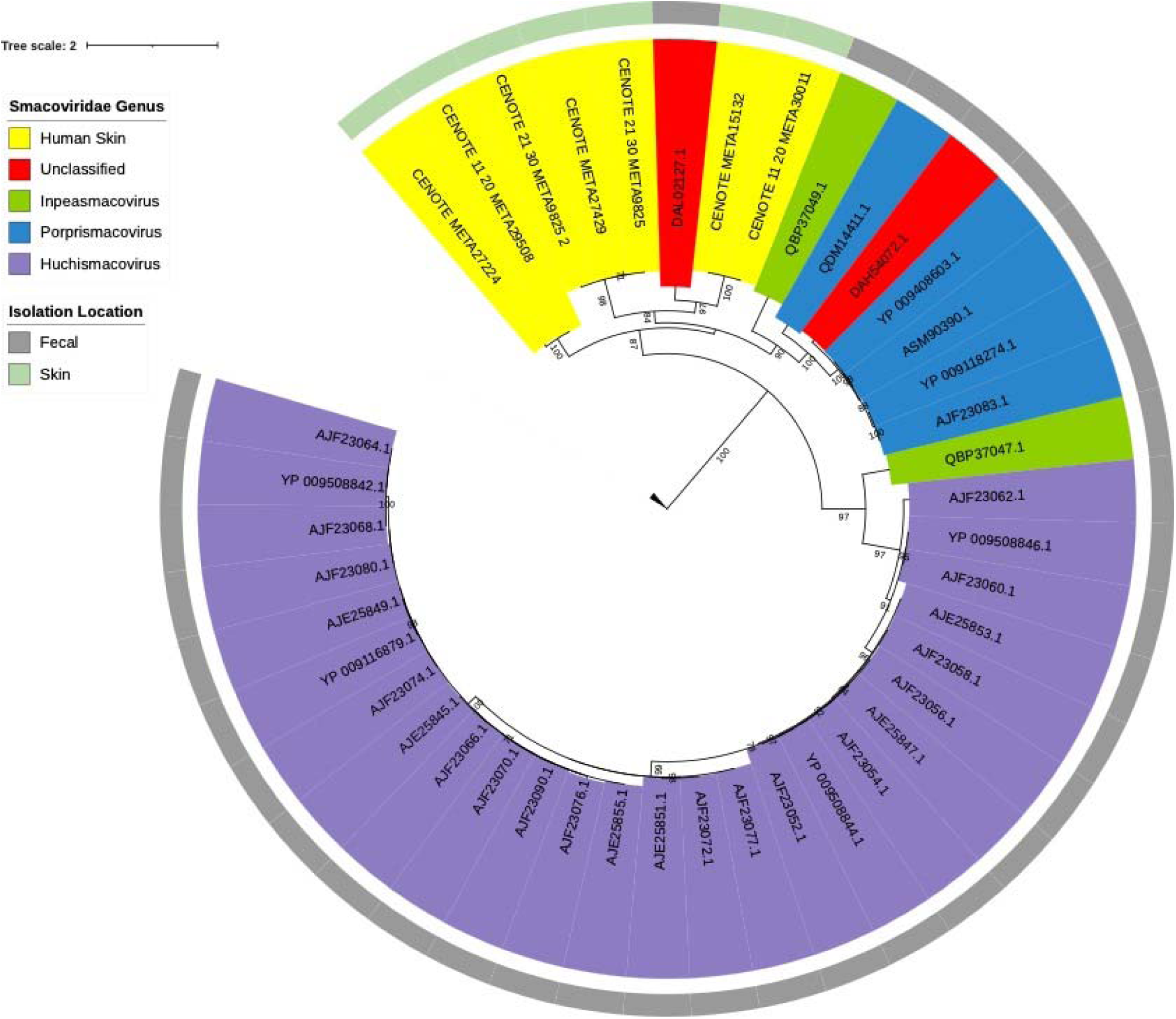
Maximum likelihood phylogenetic analysis of reference human associated *Smacoviridae* Rep protein amino acid sequences and identified novel human skin associated *Smacoviridae* Rep proteins. Reference sequences were downloaded from NCBI. Protein sequences were aligned using MAFFT, inferred using IQTree with flags for ultrafast bootstrap determination, and visualized using iTOL [59–62] using the best fit substitution model of LG+F+I+G4, as determined by IQTree. Branches with >70% bootstrap are shown. Color coordination of genus classification (clade coloring) and reported human body sampling location are represented in the figure legend. Four Rep gene amino acid sequences from differing Geminiviruses were used as an outgroup.

Though they may have shared a distant common ancestor the skin associated Rep sequences were distinctly different from that of known human associated *Smacoviridae* Rep gene sequences (Fig. 3). The closest known Rep gene reference sequence that the human skin virome database shared homology with was that of an unclassified *Smacoviridae*, which suggests that these viruses, though assumed to be *Smacoviridae* due to their family level clustering, may belong to a novel genus or genera within the *Smacoviridae*.

### Assessment of host range and phylogenetic host switching in known and human skin associated *Cressdnaviricota*

Our skin virome genome assemblies were phylogenetically distinct from that of other human associated *Smacoviridae*, therefore, we wanted to investigate if these viruses were potentially environmentally contracted and if they were more similar to that of a different host associated *Smacoviridae*. Separate maximum likelihood phylogenetic trees were constructed using the amino acid sequences of the human skin associated and all known *Smacoviridae* Rep genes (Fig. 4) and Capsid genes (Fig. 5). All *Smacoviridae* protein amino acid sequences not generated in this study were obtained from NCBI. To reduce errors in phylogenetic tree construction only full-length protein sequences were used in our analysis. When assessing Rep gene phylogeny, there was clear host conservation displayed within the tree as indicated by the clustering and small tree distances of denoted host isolation sources. However, there were many instances in which host switching could be observed throughout the tree. For the majority of the human fecal associated *Smacoviridae* (highlighted in blue), the closest instance of host switching was from that of avian, swine, and primate associated *Smacoviridae* (Fig. 4). When compared to the host switching events observed in the phylogeny of the Capsid genes there were more observed in the Rep gene phylogeny. When evaluating the Capsid gene, the closest instance of host switching was from that of bat, avian, primates and swine (Fig. 5).

**Fig 4.**
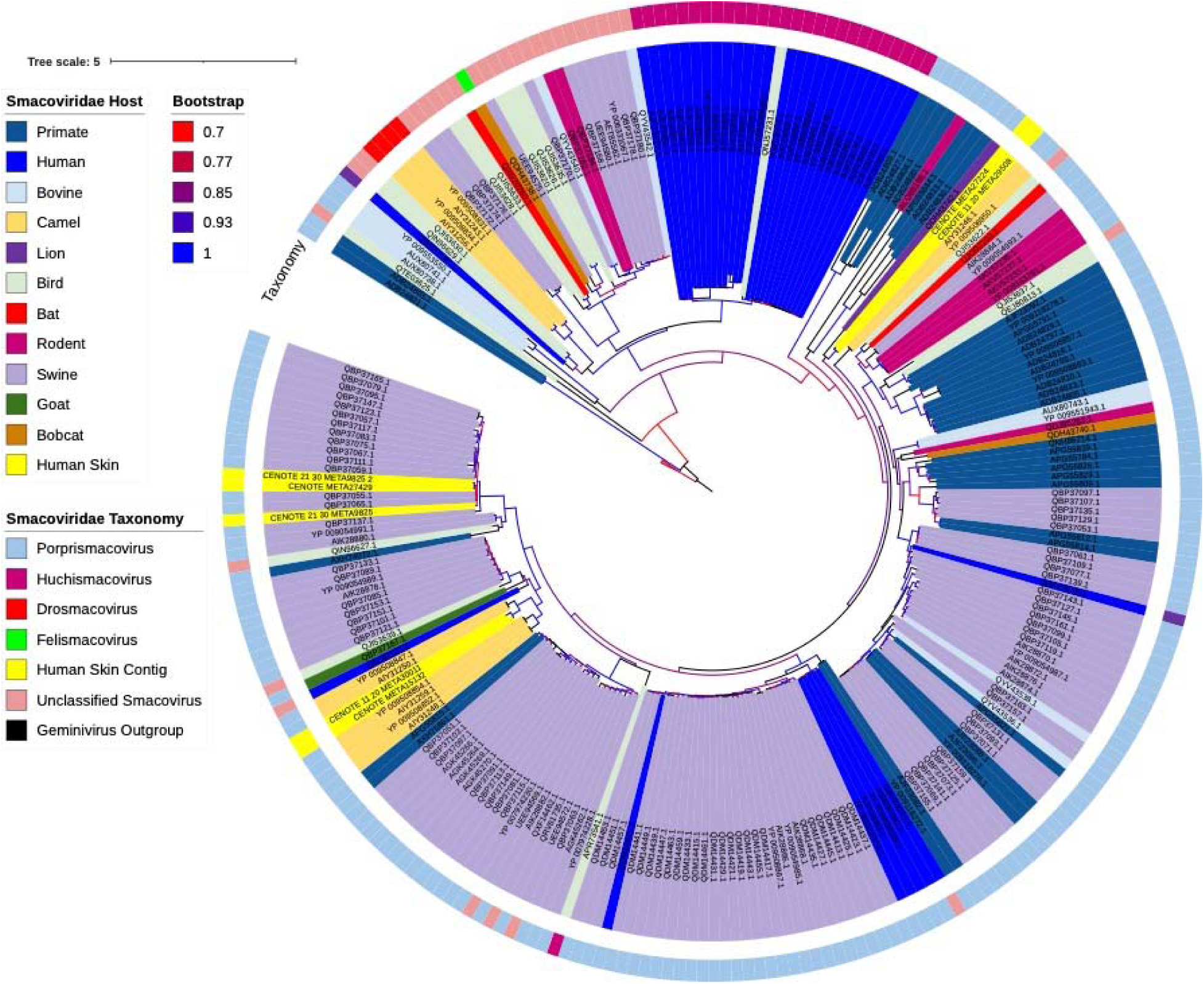
Maximum likelihood phylogenetic analysis of reference *Smacoviridae* Rep protein amino acid sequences and identified novel human skin associated *Smacoviridae* Rep proteins. Reference sequences were downloaded from NCBI. Protein sequences were aligned using MUSCLE, inferred using FastTree, and visualized using iTOL [60–62]. Branches with >70% bootstrap support are gradient colored with red representing 70% support and blue representing 100% support. Color coordination of host association and genus classification (outermost ring) are represented in the figure legend. Four Rep gene amino acid sequences from differing Geminiviruses were used as an outgroup.

**Fig 5.**
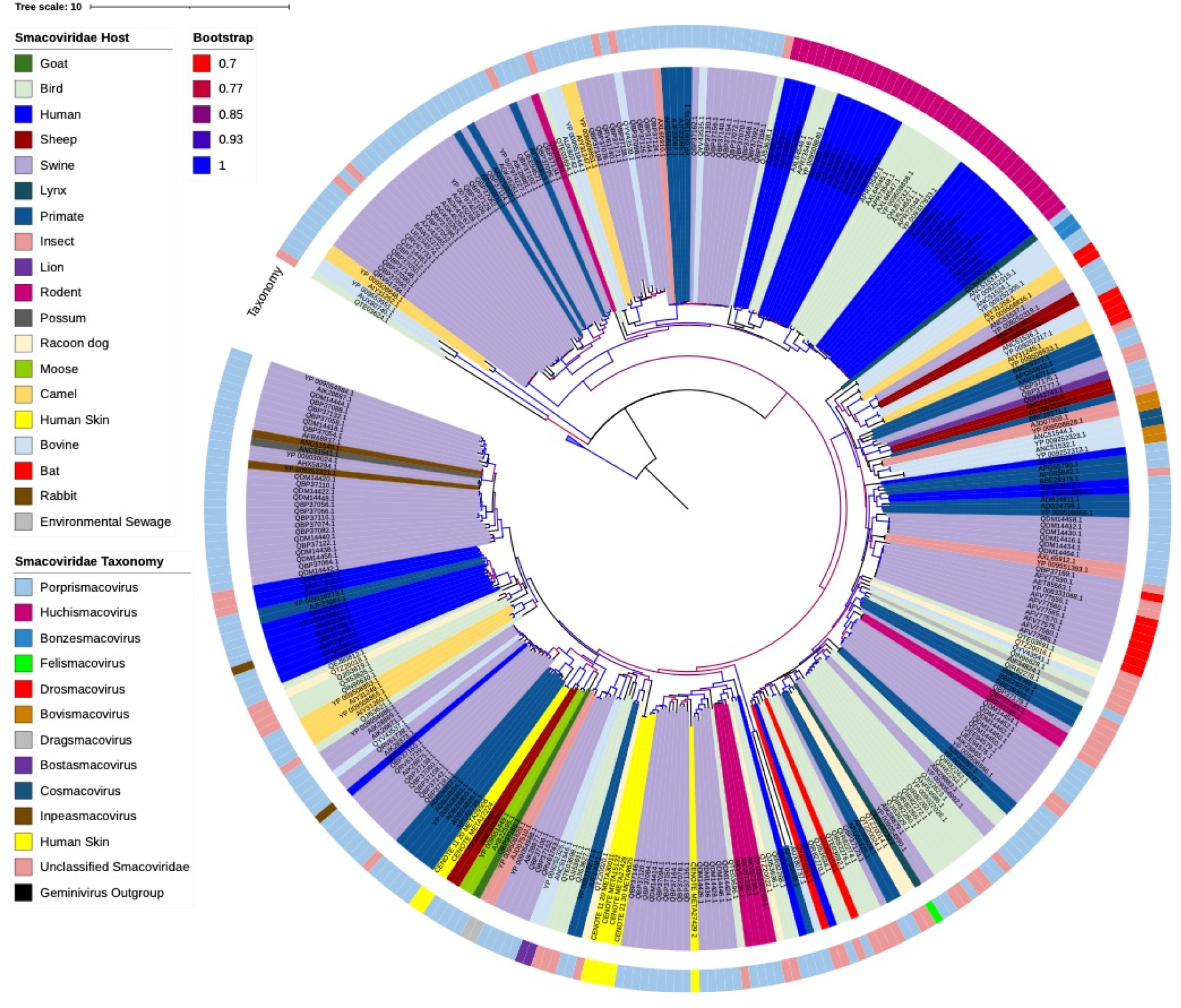
Maximum likelihood phylogenetic analysis of reference *Smacoviridae* Capsid (Cp) protein amino acid sequences and identified novel human skin associated *Smacoviridae* Cp proteins. Reference sequences were downloaded from NCBI. Protein sequences were aligned using MUSCLE, inferred using FastTree, and visualized using iTOL [60–62]. Branches with >70% bootstrap support are gradient colored with red representing 70% support and blue representing 100% support. Color coordination of host association and genus classification (outermost ring) are represented in the figure legend. Four Rep gene amino acid sequences from differing Geminiviruses were used as an outgroup.

*Genomoviridae* viral Rep genes were also investigated for host range associations and the potential for animal and/or environmental vectors for human host infection. The phylogeny of the *Genomoviridae*, which we constructed using maximum likelihood, showed immense host range and putative host switching (Fig. 6). Though there are more identified *Genomoviridae* genomes as compared to that of the *Smacoviridae*, there still was an increased amount of viral host switching observed within a phylogenetic grouping than that of *Smacoviridae*. To better visualize host range comparisons of *Smacoviridae* and *Genomoviridae* on a lower level of taxonomic grouping, we constructed sequence similarity networks to cluster genomes by genus level taxonomy (Fig. 7A, 7C). We note that some spurious or incorrectly clustered Rep genes exist within these networks, however, these instances were few. The observed ambiguity of amino acid conservation at the family and genus level is not surprising due to the genomic variability of these viruses [28].

**Fig 6.**
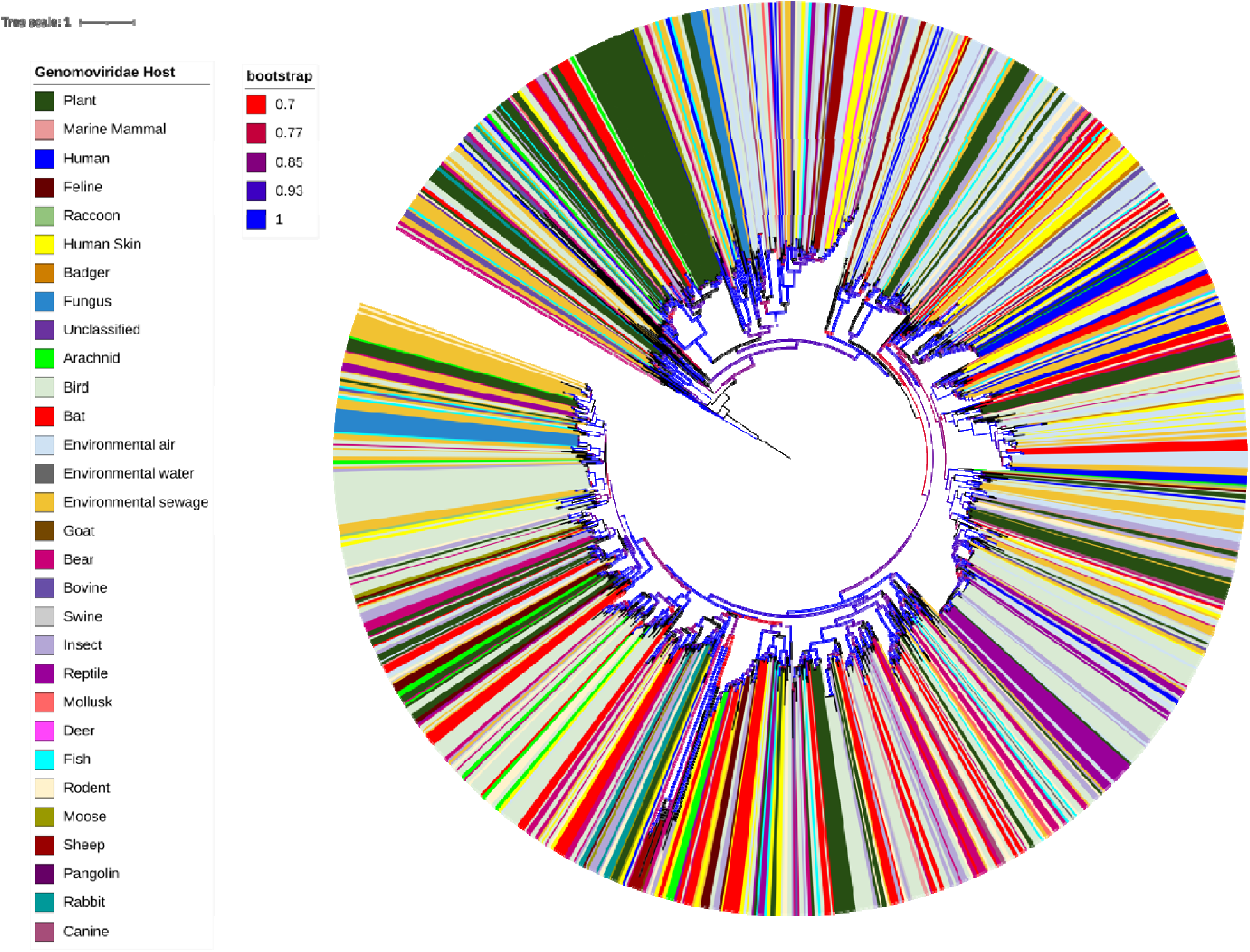
Maximum likelihood phylogenetic analysis of reference *Genomoviridae* Rep protein amino acid sequences and identified novel human skin associated *Genomoviridae* Rep proteins. Reference sequences were downloaded from NCBI. Protein sequences were aligned using MUSCLE, inferred using FastTree, and visualized using iTOL [60–62]. Branches with >70% bootstrap support are gradient colored with red representing 70% support and blue representing 100% support. Color coordination of host association are represented in the figure legend. Four Rep gene amino acid sequences from differing Geminiviruses were used as an outgroup.

**Fig 7.**
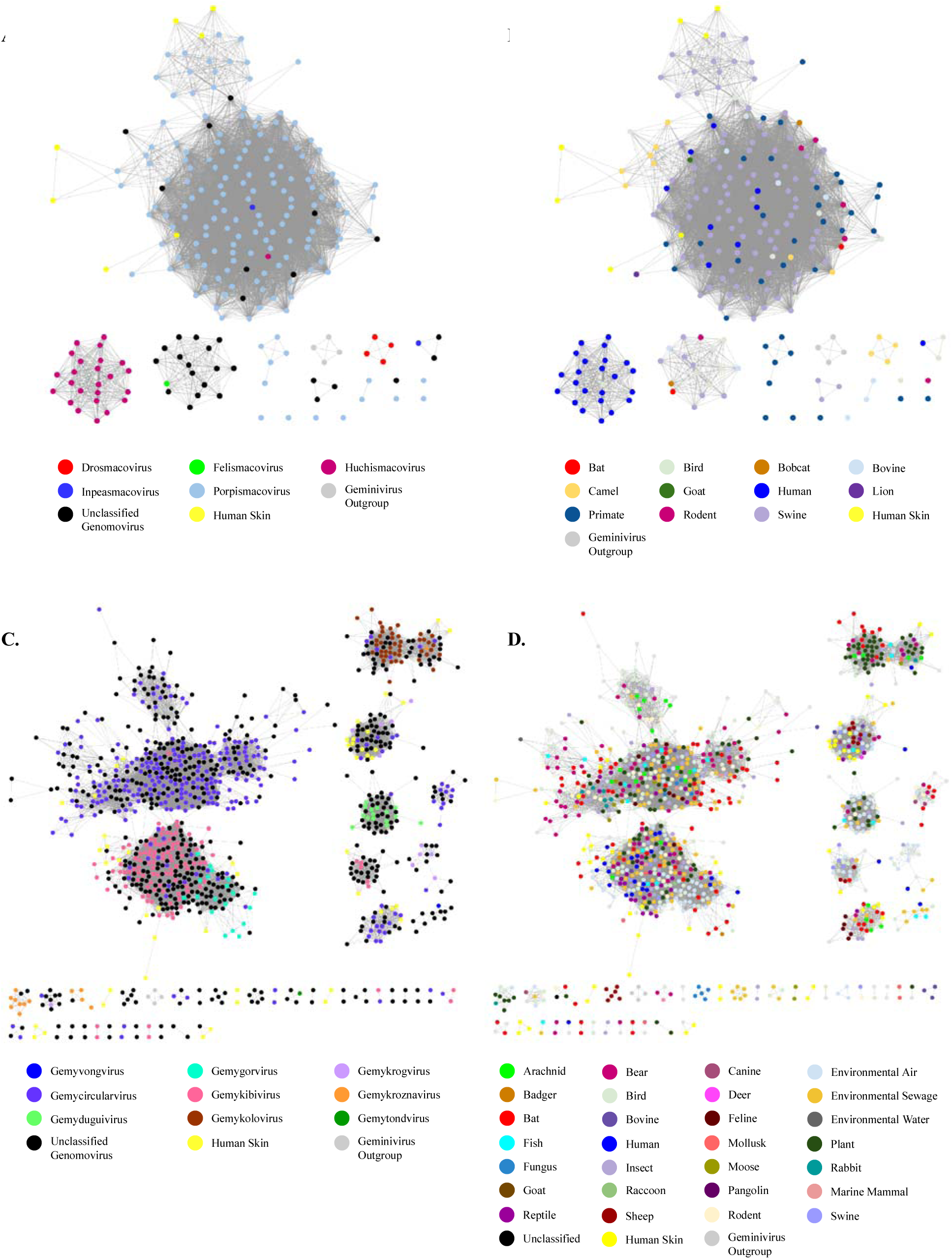
Sequence similarity networks of *Smacoviridae* and *Genomoviridae* reference sequences obtained from NCBI and the human skin associated Smacoviruses identified in this study. EFI-EST was used to conduct pairwise alignments of the conserved amino acid sequences of the Rep gene [55]. For Genus level clustering an E-value cutoff of 10^-60^ was used for *Smacoviridae* and 10^-100^ for *Genomoviridae*. Cytoscape was used for network visualization and color labeling [56]. (A.) Color designations represent taxonomy classifications and (B.) host associations as represented in the figure legends. Human skin associated Smacoviruses identified in this study are highlighted in yellow. Four Rep gene amino acid sequences from differing Geminiviruses were used as an outgroup control for this network clustering. (C.) Color designations represent taxonomy classifications and (D.) host associations as represented in the figure legends. Human skin associated Genomoviruses identified in this study are highlighted in yellow. Four Rep gene amino acid sequences from differing Geminiviruses were used as an outgroup control for this network clustering.

When evaluating these networks, human skin viruses (highlighted in yellow in all figures) clustered similarly to what was shown in the phylogenetic analysis using maximum likelihood estimations (Fig. 4, 6). These sequence similarity networks displayed distinct groupings that can be considered that of hypothetical genus level groupings (Fig. 7). For some novel genus level groupings identified in this study, we observed that host associations were conserved within specific hypothetical genus level designations (Fig. 7D). However, for all of these instances there are small numbers of genomic samples present in these groupings. In comparison to the networks of the *Smacoviridae* hypothetical genus level taxonomic groupings (Fig 7A-B), host specificity recognized at the genus level was more conserved in that of the *Smacoviridae* than that of the *Genomoviridae* (Fig. 7C-D).

## DISCUSSION

There is a substantial lack of knowledge about the viruses associated with the human skin as well as the potential for those viruses to spread via contact and impact human health. Most human skin virome studies have been directed at phage population dynamics [5,6,8], known opportunistic skin pathogens such as Papillomaviruses and Polyomaviruses [4,38,39], and dsDNA viral populations [5,6,8]. Our previous study [9] identified highly abundant stable ssDNA viruses, however, the classification of these viral metagenome assemblies was ambiguous due to poor representation in reference databases. Our goal in this study was to address the gap of knowledge in the breadth of known and novel viral ssDNA viruses on the human skin to improve current viral reference databases and to investigate potential novel human viral pathogens.

We identified 421 unique ssDNA viral genome assemblies from human skin virome samples collected from 60 individuals spanning three anatomical locations sampled over a six-month time-period. Although many of these viral assemblies had low sequence similarity to publicly available viral genomes, which highlights the lack of knowledge of human skin ssDNA viral populations, many of these novel genome assemblies exhibited similar genomic composition to the *Cressdnaviricota*. Of the ssDNA viral genomes identified, the majority (∼65%) were that of the Rep-gene containing *Cressdnaviricota*. No previous studies have sought to exclusively address the *Cressdnaviricota* populations on the human skin. These viruses are of interest to human health due their association with systemic pathogenesis in animals such as swine and Porcine Circovirus 2 (PCV2) or bird beak and feather disease [40, 41], as well as associations with acute respiratory and encephalitis cases in humans [23,29,30,37].

The viral genomes we identified in this study contain Rep genes that are clustered predominantly with known eukaryotic infecting *Genomoviridae* (Fig. 1E). A few other assembled genomes clustered with *Circoviridae* and the human and animal fecal associated family *Smacoviridae*. Putative *Circoviridae* genomes containing a Rep gene derived from our human skin samples clustered separately from the main *Circoviridae* cluster derived from sequences deposited to public databases. We did not find any human skin associated viral genomes that clustered with plant-infecting *Nanoviridae* and *Geminiviridae*. We did identify distinct novel clusters exhibiting family-level sequence similarity from our dataset. These distinct clusterings represent putative novel families of *Cressdnaviricota* or belong to novel *Cressdnaviricota* families such as CressV1 [20], however, further phylogenetic analysis with increased sampling is needed to confirm these new family level groupings.

Due to the unique genomic nature of the *Cressdnaviricota*, which are characterized by containing only two genes, a replicase (Rep) and Capsid gene, as well as a lack of host-specific virulence genes, it is hypothesized that the *Cressdnaviricota* have the potential to be pathogenic to any host cell they are able to enter due to the replication efficiency and expression of their own rolling cycle-replicase (Rep) genes [42]. However, understanding of overall host range and cell specificity of specific families within the *Cressdnaviricota* is still in its infancy [19]. Previous studies [19,43,44] have set out to address the host specificity of the families within the *Cressdnaviricota* and to hypothesize that they have the ability to switch between hosts via Capsid mutations and recombinational events, similar to what has been recently observed in unrelated Coronaviruses [45, 46].

There are two families with notable agricultural importance within the *Cressdnaviricota* – the *Geminiviridae*, which are pathogenic on a broad host range of plants, and the *Circoviridae*, which are associated with systemic pathogenesis in animals such as “post-weaning stress syndrome” in swine [41] and “bird beak and feather disease” in fowl [40]. However, for other families within the *Cressdnaviricota* little is known about their ecology, host specificity, and pathology. Interestingly, the two viral families where human skin associated *Cressdnaviricota* were phylogenetically associated with each other were that of *Genomoviridae* and *Smacoviridae*. Host associations and the potential for zoonotic transfer of these two families within the *Cressdnaviricota* are not understood. Without clear visible pathogenesis, it is difficult to assess if the *Cressdnaviricota* identified in our present study are typically environmentally contracted or if they are active viral infections of the human skin.

Here we showed that the family *Genomoviridae* were highly abundant, stable over time, and consistently present across the five time points from multiple skin anatomical locations assessed for the 60 study participants (Fig. 1C-E). This is consistent with our findings from our previous study where the viral family *Genomoviridae* was shown to be stable over time for 42 study participants [9]. Not only did we previously show that these viruses were a part of the core human skin virome, but when performing a phylogenetic analysis of all non-redundant *Genomoviridae* protein sequences in this study, it became clear that the *Genomoviridae* not only have broad host range but are also categorized as having associations with multiple cell types even within the same host. Due to their stability over time, we hypothesize that these viruses are established viruses that are continually replicating or newly colonizing cells from the environment, though their pathogenesis is currently unknown. This is in opposition to the *Smacoviridae* which we observe are temporal and appear to be environmentally contracted or are pathogenic in non-human hosts. Here we confirm the persistent presence of the *Genomoviridae* on the human skin and conclude there is a high probability for transfer, and, due to their immense host range, there is potential for host-switching events to occur through contact. We therefore hypothesize there is potential for zoonotic infections via host-switching from the *Genomoviridae*.

We found many instances of clustering of unclassified *Genomoviridae* Rep amino acid sequences, and we hypothesize that these may be novel families and genera of the *Genomoviridae*. Additionally, clustering of the unclassified *Genomoviridae* offers insight and resolution in the classification of these novel viral genus and species level classifications. Though we were able to cluster at the level of viral genus on the basis classification, when highlighted based on host, it was evident that even within genus level classification there is an immense host range.

Of interest, we found six human skin associated viral genomes that contained a Rep gene sequence (and one partial Rep sequence) and that clustered with the fecal associated *Cressdnaviricota* family, *Smacoviridae*. The identification of *Smacoviridae* related viruses is of human health interest since an isolate of the *Smacoviridae* has been implicated with cases of severe encephalitis from patients in France and gastrointestinal disease in a cohort of children from South America [30,37,43]. However, this finding was not expected as the *Smacoviridae* have not previously been associated with, isolated from, or been found within or on, animal or human tissue samples. The presence of novel *Smacoviridae* on human skin does not imply active infection of these viruses in human skin cells, especially since it has been hypothesized that this viral family broadly infects Archaea [27]. However, the identification of this viral family is of importance because very little is known about the Archaeal population on the skin and the potential health implications of shifts in Archaeal populations on the human skin through viral pathogenesis. Though abundantly present on the skin of several study participants, the *Smacoviridae* were not persistently present on any individual’s skin over time. Due to the combination of high abundance when present and their transient nature across the sampling time frame, we believe that the *Smacoviridae* is environmentally deposited on the human skin and not a part of the core human skin virome. Evaluations of the capsid gene protein sequences revealed the closest non-human host which exhibited the presence of the *Smacoviridae* was that of sheep, raccoon dog, and swine (Fig. 4), although the phylogenetic distance of the tree constructed with both the Rep and Capsid genes was not particularly close suggesting the presence of an intermediary vector, or perhaps evolutionary time had passed between host-switching events. When comparing just the Rep genes, the closest non-human host associated with the *Smacoviridae* was that of a camel, swine, and a lion (Fig. 5). The incongruity in closest non-human host between the Rep and Capsid genes, as well as the phylogenetic differences from viral genomes isolated from human fecal and human skin, gives credence to a large host range of these viruses perhaps again highlighting the hypothesis that they are hosts of Archaea. Their associations with human disease are an anomaly and warrant further investigation.

Due to the high recombination rates and the subsequent difficulty in Rep gene alignment, we were not surprised to see multiple clusters of the *Circoviridae* (Fig. 1E) as has also been observed in other studies [19]. However, due to the low number of reference *Circoviridae* Rep sequences in these clusters and the potential for classification errors from public databases we cannot say definitively that the novel human skin associated *Circoviridae*-like genomes from this study should in fact be identified as *Circoviridae*. We therefore did no further analysis on the *Circoviridae*-like genomes past their relative abundance and persistence throughout our dataset. The unknown diversity of the Circoviridae-like genomes warrants further dedicated study.

When considering the Capsid gene phylogeny with host mapped across the phylogenetic tree, it is evident that host switching occurs frequently indicating greater instances of recombination events than that of the Rep gene. The Capsid gene is crucial for recognition in host-specificity and therefore it is not surprising to see greater genetic variation or instances of recombination events within this gene as compared to that of the Rep gene which has been shown to be more conserved across *Cressdnaviricota* families [28].

Because the human skin is our first contact with the environment, it plays a crucial role in restricting viral and microbial pathogen spread and transmission, therefore, further studies focused on the transfer, contraction, pathogenesis, and potential animal vectors of this viral family are warranted. Most research has focused on RNA viruses as emerging pathogens and zoonotic and host vector prediction with very little known about DNA viruses. Due to the extreme host range, adaptive capabilities, and presence of these viruses on the human skin we believe that the findings of this research warrant further investigation into prediction of host vectors, emergence of novel *Cressdnaviruses*, and zoonotic transfer of these small ssDNA viruses, especially the instances of animal pathogenic versions of these viruses. Findings from this study underscore the need for more ssDNA viral focused studies into prediction of host vectors, emergence of novel strains, and zoonotic transfer, especially for instances of animal pathogenic versions of these viruses.

## MATERIALS AND METHODS

### Sample Collection of the Viral DNA Component of Human Skin

Human skin viral metagenomic samples were collected from 60 individuals who were not on an antibiotic regime. At each time of collection, the participants of the study were sampled at three anatomical locations of their skin – left hand, right hand, and scalp – and were individually swabbed using a dual dry and 1xPBS wet swabbing method as described in Graham et al. [9]. A time-series of sampling was performed across a longitudinal six-month period to represent the initial sampling (day 0), and two-week, one-month, three-month, and six-month time intervals from the initial collection date. Negative control swabs saturated in 1xPBS were collected at each sampling to evaluate potential environmental contaminants. All negative and blank controls collected were processed and sequenced to evaluate environmental and lab contamination. Post collection, skin swabs were stored at -20°C until further processing.

Cellular material was purified from swabs by saturating the swabs with 200ul of 0.02µm filtered sterile 1xPBS and placing them in a CW Spin Basket (Promega, Madison, WI, USA). The saturated swabs were then centrifuged at 16,000xg for 10 min to elute viral and cellular material from the swab. The filtrate was then passed through a 0.22µm filter to enrich for viral particles and remove cellular and bacterial contaminants. Sample DNA was extracted from the viral enriched filtrate using the QiAmp Ultra-Sensitive Virus Kit (Qiagen, Hilden, Germany) and was subjected to whole genome amplification (WGA) using multiple displacement amplification (MDA) implemented with the TruePrime WGA Kit (Syngen Biotechnology, Inc, Taipei City, Taiwan) following standard product protocols. Following WGA, the samples were quantified using the DeNovix dsDNA High Sensitivity Kit using the DeNovix DS-11 Spectrophotometer/Fluorometer (DeNovix, Inc, Wilmington, DE, USA).

### Metagenomic Sequencing of Skin Samples

For each sample, 100 nanograms of amplified DNA was sheared using sonication, implemented with a Bioruptor (Diagenode, Denville, NJ, USA) with three cycles of 30s on and 90s off as per manufacturer instructions, to obtain a mean DNA length distribution of 600bp. The sheared DNA was prepared for sequencing library preparation with the NEBNext Ultra II Library preparation kit (New England Biolabs, Ipswich, MA, USA) according to the manufacturer’s protocol. During the sequencing library construction, we used the recommended approximate insert size of 500 to 700 bp for the adapter-ligated DNA, followed by PCR enrichment of five cycles of denaturation and annealing/extension. The base-pair size distribution of the resulting sequencing library was quality checked using the high sensitivity chip on an Agilent 2100 Bioanalyzer (Agilent Technologies, Inc, Santa Clara, CA, USA). Prior to sequencing, libraries were quantified using the DeNovix dsDNA High Sensitivity Kit (DeNovix, Inc, Wilmington, DE, USA) and diluted to consistent concentrations before being barcoded and subsequently pooled. A single sequencing library was then sequenced on a Illumina HiSeq 2500 platform machine (Illumina, Inc, San Diego, CA, USA) using a 150 bp paired-end sequencing strategy. All raw sequencing data produced for this study has been deposited in the NCBI Short Read Archive (SRA) under the accession code PRJNA754140.

### Assembly of Metagenomic Sequencing Data

The sequencing data we generated was first evaluated for quality using FastQC v.0.11.9 [47] and trimming was performed to remove low quality reads with a filter threshold of Q30 and a minimum length threshold of 75 bp using Sickle v.1.3.3. [48]. PhiX sequencing contamination was removed using the BBDuk command with standard operational flags for PhiX removal as recommended by the BBMap suite of tools [49]. Following quality filtering, human genome contamination was removed to mitigate host sequence contamination by mapping all reads to the human genome (hg19) using BBMap with standard operational flags [49] for high precision mapping with low sensitivity to lower the risk of false positive mapping.

Using MEGAHIT v.1.2.8 [50], filtered reads were assembled using two approaches, 1) a master meta-assembly using all reads, and 2) assembly within each sample. Assembled genomes were quality assessed using QUAST v.5.0.2 [51]. Subsequent contigs greater than 1000bp were retained for all downstream analysis. The same bioinformatic assessment of samples was performed on the negative control samples of the sequencing reads. A meta-assembly was constructed using all negative control reads and the sample genome assemblies were mapped to the negative control contigs greater than 1000bp using BWA-mem [52] to remove reads resulting from possible contamination. Unmapped viral contigs greater than 1kb were retained for further analysis.

### Identification and Annotation of Circular ssDNA Viral Contigs

Circular viral contigs were identified and annotated using Cenote-Taker2 [53] with condition flags to filter out plasmids and recommended conditions for whole genome shotgun metagenomic assembly data. Identified putative circular viral contigs were then manually separated into nine groups (Anellovirus, *Cressdnaviricota*, *Inoviridae*, Papillomavirus, Polyomavirus, Microvirus, Unclassified phage, and partial unclassified genomes containing only a capsid gene) based on their protein annotation and classification outputs from Cenote-Taker2.

### Public Data Mining

To incorporate known *Cressdnaviricota* from publicly available databases, we queried amino acid sequences from NCBI, with a known family level classification of the *Cressdnaviricota*, for both the Rep and Capsid gene sequences. Sequences obtained from NCBI were preliminarily aligned, inspected by eye, and any partial or truncated protein sequences were removed from the analysis.

### Sequence Similarity Network Construction

To classify the novel Rep containing skin associated viruses a data set containing all Rep *Cressdnaviricota* sequences with known family level classification obtained from NCBI was constructed. This dataset was then further reduced by clustering sequences at a 90% amino acid similarity using CD-HIT [54]. We used EFI-EST to construct a sequence similarity network using pairwise alignments of the 90% similar Rep gene and the human skin associated Rep gene amino acid sequences with an E-value cut off of 10^-60^ [19, 55]. Additional networks were constructed using all full-length available Rep amino acid sequences for the viral families *Smacoviridae* and *Genomoviridae* paired with the corresponding novel human skin associated Smacoviruses and Genomoviruses identified in this study, respectively. We employed an E-value cut-off of 10^-60^ for the *Smacoviridae* network and 10^-100^ for the *Genomoviridae* network to represent taxonomic clustering at the viral genus level. All networks were visualized, and color annotated using Cytoscape v.3.8.2 [56].

### Phylogenetic Analysis

Protein amino acid sequence alignments were performed using MAFFT v.7.407 [57] with optimization for accurate local alignment (using the LNSI model). Phylogeny was then inferred using IQ-TREE v.1.6.7 with conditions for ultrafast bootstrap determination [58, 59]. For larger trees containing more than 300 protein amino acid sequences, alignments were performed using MUSCLE v.3.8.1551 [60]. Phylogeny of the larger alignments was inferred using FastTree v.2.1.11 [61]. Outgroups consisting of four Geminivirus strains of the sweet potato curl virus Rep protein (ACY79444.1, ACY79438.1, ACY79480.1, ACY79486.1) or capsid protein (ACS44338.1, ABS83269.1, ALK03540.1, ALK03534.1) amino acid sequences were used for all trees. Host specification and annotation files were constructed using information provided from the NCBI protein database and edited manually for consistency. Trees were visualized and edited using iTOL v.6.5.2 [62].

### Mapping of Sequencing Reads to Cressdnavirus Contigs and Annotated Proteins

A reference database was constructed containing all Cenote-Taker2 identified Rep genes containing genome assemblies and all annotated Rep and Capsid gene nucleotide sequences identified from our human skin dataset. Quality filtered and trimmed sequencing reads were mapped to the reference database sequences using Bowtie2 v.2.3.5. [63]. Subsequent mapped read counts were acquired and filtered using SAMtools [64].

### Evaluation of *Smacoviridae* Virus Abundance, Stability, and Diversity Across Subjects

We then used data in the form of the mapped read counts, collected sampling metadata, and family level sequence similarity clustering classifications to assess viral persistence over time (i.e., stability) and abundance within the study population using R v.3.6.3 [65]. Persistence over time was calculated by using binary presence vs absence metrics for each family grouping for each study participant by anatomical skin site locations across the five time point collections. Abundance was calculated by summing across the five time points for each for the family level groupings for each anatomical location within a subject. For abundance and persistence comparisons, only subject sample comparisons where four out of the five time points or more were collected for each skin site location were used. Results bar graph and heatmaps were visualized using the R packages ggplot2 v.3.3.3 and Pheatmap v.1.0.12 in R v.3.6.3 [65, 66].

## ACKNOWLEDGEMENTS

We thank all study participants for their contribution to this study. Additionally, we would like to thank W. Tom, C. Anderson, and other individuals in the Fernando Lab who assisted with experimentation and bioinformatic support. This work was completed using the Holland Computing Center (HCC) at the University of Nebraska, which receives support from the Nebraska Research Initiative.

## DATA AVAILABILITY

Raw sequencing data is available through the NCBI Short Read Archive (SRA) under the project accession code PRJNA754140. Project metadata and bioinformatic pipeline scripts described in this manuscript are publicly available at: https://github.com/HerrLab/Graham_2022_ssDNA_Virome_Cress.

## AUTHOR CONTRIBUTIONS

EG, PA, SF, and JH contributed to the experimental design and conceptualization of this study; MA was responsible for participant recruitment and sample collection; EG and SF developed virome processing methodology and prepared samples for sequencing; EG and JH developed the bioinformatic pipeline and conducted phylogenetic analysis on sequence data; JC contributed statistical support; EG and JH drafted this manuscript with input from all authors.

